# Biochemical and Genetic Evidence Supports Fyv6 as a Second-Step Splicing Factor in *Saccharomyces cerevisiae*

**DOI:** 10.1101/2023.01.30.526368

**Authors:** Karli A. Lipinski, Katherine A. Senn, Natalie J. Zeps, Aaron A. Hoskins

**Affiliations:** Department of Chemistry, University of Wisconsin-Madison, Madison, WI 53706; Department of Biochemistry, University of Wisconsin-Madison, Madison, WI 53706

**Keywords:** Fyv6, FAM192A, spliceosome, yeast, RNA splicing

## Abstract

Precursor mRNA (pre-mRNA) splicing is an essential process for gene expression in eukaryotes catalyzed by the spliceosome in two transesterification steps. The spliceosome is a large, highly dynamic complex composed of 5 small nuclear RNAs and dozens of proteins, some of which are needed throughout the splicing reaction while others only act during specific stages. The human protein FAM192A was recently proposed to be a splicing factor that functions during the second transesterification step, exon ligation, based on analysis of cryo-electron microscopy (cryo-EM) density. It was also proposed that Fyv6 might be the functional *S. cerevisiae* homolog of FAM192A; however, no biochemical or genetic data has been reported to support this hypothesis. Herein, we show that Fyv6 is a splicing factor and acts during exon ligation. Deletion of *FYV6* results in genetic interactions with the essential splicing factors Prp8, Prp16, and Prp22; decreases splicing *in vivo* of reporter genes harboring intron substitutions that limit the rate of exon ligation; and changes 3’ splice site (SS) selection. Together, these data suggest that Fyv6 is a component of the spliceosome and the potential functional and structural homolog of human FAM192A.

## INTRODUCTION

The removal of introns from pre-mRNA molecules is carried out by the spliceosome, a large macromolecular complex made up of five small nuclear RNAs (snRNAs) and dozens of proteins which assemble de novo on each pre-mRNA substrate. Splicing consists of two, stepwise transesterification reactions in which the 5’ splice site (the boundary between the 5’ exon and the intron; 5’ SS) is first cleaved by formation of a lariat intron and then the intron is released concomitant with exon ligation by attack of the 5’ exon at the 3’ SS. Spliceosome composition changes dramatically throughout the course of splicing due to the sequential arrivals and departures of different components as well as large-scale conformational changes (Plaschka, et al., 2019). This results in the formation of several intermediate complexes with distinct architectures during the reaction, many of which have now been visualized by cryo-electron microscopy (cryo-EM) (Plaschka et al., 2019). Cleavage of the 5’ SS is completed during the transition from the spliceosome B* to the C complex, and exon ligation occurs during the transition between the C* and P (product) complexes. While some components of the spliceosome remain part of the machine throughout the reaction, others transiently associate, dissociate, or re-arrange to interact with the catalytic site only at specific times. Just prior to 5’ SS cleavage, the 1^st^ step factors (Cwc25, Isy1, and Yju2) function to juxtapose the 5’ SS and branch site (BS) (Wan et al., 2019; Liu Y-C et al., 2007; Villa and Guthrie, 2005; Chiu et al., 2009; Wilkinson et al., 2021). Cwc25, Isy1, and Yju2 are then released after 5’ SS cleavage, and 2^nd^ step factors bind (Slu7, Prp18) or are repositioned (Prp17) to facilitate exon ligation (Plaschka et al., 2019; James et al., 2002; Yan et al., 2017; Fica et al., 2017; Ohrt et al., 2013; Tseng et al. 2011). Proper progression through splicing requires the coordinated association and dissociation of these 1^st^ and 2^nd^ step factors with the active site and these transitions are enabled, in part, by ATP-dependent DExD/H-box helicases. The ATPase Prp16 promotes rearrangement of the spliceosome active site and splicing factor release between the 1^st^ and 2^nd^ step of splicing (Schwer and Guthrie, 1992; Semlow et al., 2016), while Prp22 promotes release of the mRNA product from the spliceosome after exon ligation (Wagner et al., 1998; Schwer, 2008).

Recently, a putative new 2^nd^ step factor (FAM192A or PIP30) was identified for the human spliceosome. The protein was found by fitting unassigned density present in cryo-EM maps of spliceosomes transitioning between conformations competent for the 1^st^ and 2^nd^ steps (Zhan et al., 2022). Depletion of FAM192A from human nuclear extracts reduced *in vitro* splicing but adding purified protein back did not restore this activity, potentially due to simultaneous depletion of other, unidentified splicing factors (Zhan et al., 2022). Consequently, its role in splicing is still poorly defined.

Interestingly, Zhan et al. also identified a potential FAM192A homolog, Fyv6 (Function required for yeast viability 6), in *Saccharomyces cerevisiae* (hereafter, yeast) despite having less than 20% sequence identity (**Fig. 1A**). (It should be noted, however, that this level of sequence identity is similar to that between yeast and human homologs of the other 2^nd^-step factors Slu7 and Prp18). The predicted AlphaFold structure of Fyv6 (Jumper et al., 2021) was able to be modeled into unassigned EM density from yeast C* spliceosome complexes (previously labeled as unknown protein X) (Zhan et al., 2022) (**Fig. 1B**). Prior to this work, Fyv6 had been detected by mass spectrometry analysis of purified B^act^ and C complex spliceosomes (Warkocki et al., 2009; Fabrizio et al., 2009) as well as postulated to be responsible for unassigned density in a cryo-EM structure of a yeast P complex spliceosome (referred to as UNK in that structure, **Fig. 1C**) (Liu et al., 2017). In both the C* and P complex spliceosomes, the unassigned EM density is located in a position that could significantly impact splicing chemistry: in C* the density contacts core splicing factors Cef1, Syf1, and Prp8 while in P complex it contacts these factors in addition to the lariat intron, Slu7, and Prp22 (**Figs. 1**, **S1**). Together the combined cryo-EM and mass spectrometry data hint at Fyv6 functioning during splicing; however, no genetic or biochemical evidence for this has been reported.

**Figure 1.**
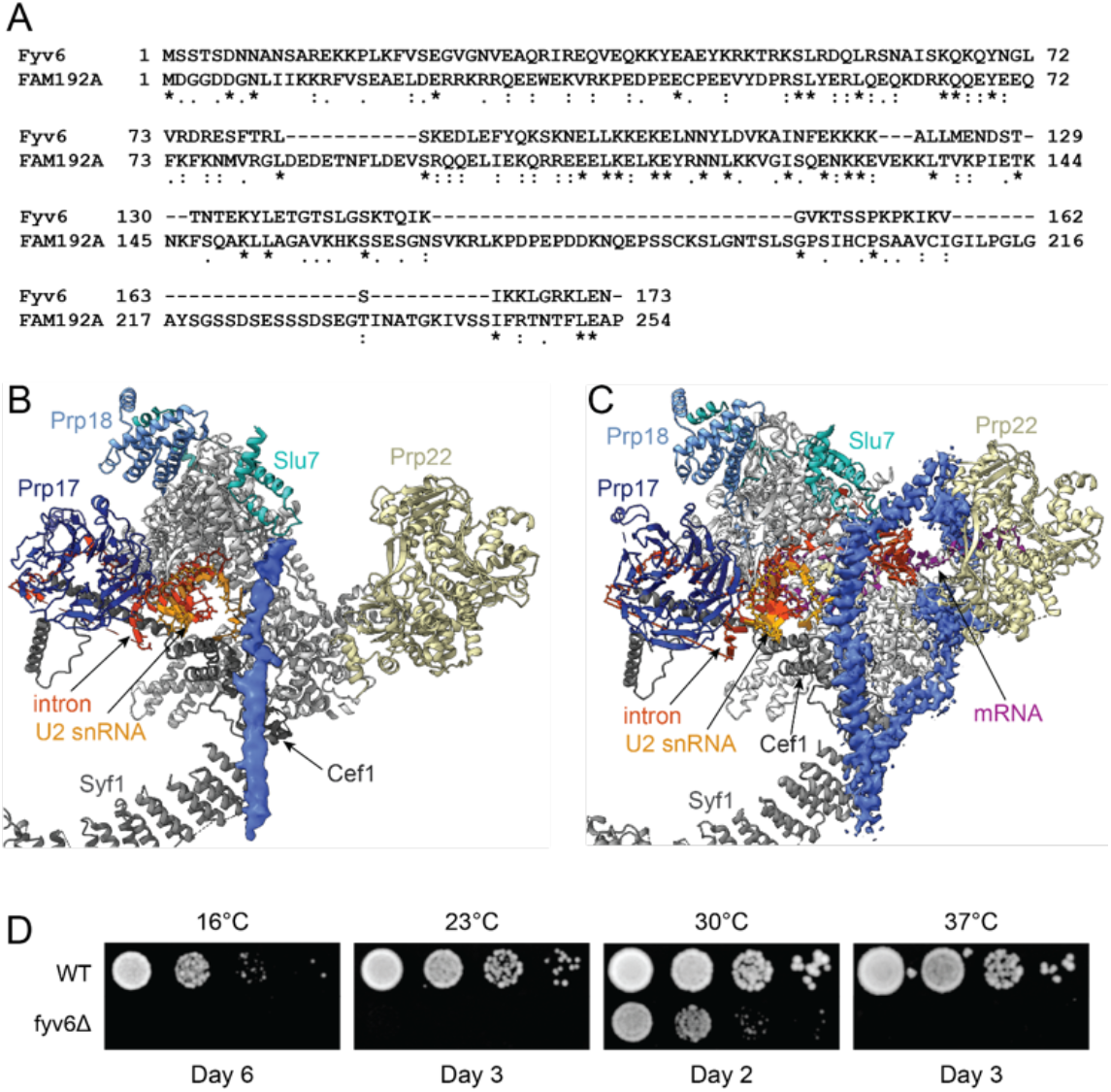
Sequence Alignment of Fyv6 with FAM192 and Unassigned EM Density in Yeast Spliceosome Structures. **A)** Sequence based alignment of *S. cerevisiae* Fyv6 and human FAM192A using EMBOSS Needle (Needleman and Wunsch, 1970). **B), C)** Superposition of the atomic models for the spliceosome C* (panel B, 5MQ0) and P (panel C, 6BK8) complexes with the unassigned EM density shown in blue spacefill. Images were prepared using ChimeraX (Pettersen et al., 2021). **D)** Impact of *fyv6D* on yeast growth at various temperatures. Plates were imaged on the days shown.

Fyv6 is a poorly studied and nonessential yeast protein originally identified in a screen for mutants sensitive to K1 killer toxin (Pagé et al., 2003). Since identification, Fyv6 has appeared in genetic screens for mutants with sensitivity to heat (Auesukaree et al., 2002), calcineurin inhibitor FK506 (Viladevall et al., 2004), and changes to cell size (Maitra et al., 2019). Fyv6 is localized to the nucleus and has previously been proposed to play a role in nonhomologous end joining, but little is known about its function or interacting partners (Huh et al., 2003; Wilson, 2002). Here, we studied the function of Fyv6 during splicing by probing genetic interactions between Fyv6 and the splicing factors Prp8, Prp16, and Prp22. In addition, we assayed its impact on splicing *in vivo* by deleting *FYV6* and with use of ACT1-CUP1 splicing reporters. Together, these data are consistent with Fyv6 functioning as a 2^nd^-step splicing factor in yeast.

## RESULTS AND DISCUSSION

### Genetic Interactions between Fyv6 and Prp8 1^st^ or 2^nd^-Step Alleles

To examine a potential role for Fyv6 in splicing, we deleted *FYV6* from the yeast genome, confirmed the deletion by PCR (**Fig. S2**), and assayed for genetic interactions with known splicing factors. While *FYV6* is nonessential for yeast viability, its deletion does cause a slow growth defect at 30°C and both cold and temperature sensitivity (*cs* and *ts*) phenotypes at other temperatures (**Fig 1D**). We first tested for genetic interactions with the essential spliceosome component Prp8. Prp8 is a central protein in the spliceosome that scaffolds the active site RNAs and can impact equilibria between the intron branching and exon ligation reactions through structural rearrangement (Query and Konarska, 2004; Fica and Nagai, 2017). As such, multiple alleles of Prp8 stabilize the spliceosome in either the 1^st^ or 2^nd^-step conformation at the expense of the other state (**Fig. 2A**) (Umen and Guthrie, 1995a; Schneider et al., 2004; Query and Konarska, 2004; Liu L. et al., 2007). Moreover, alleles of 2^nd^ step factors Prp18 and Slu7 (*prp18-1, slu7-1, slu7-ccss*) are synthetically lethal with a 1^st^ step allele of Prp8 (*prp8-101* or Prp8^E1960K^; Umen and Guthrie, 1995b), presumably since both alleles work in concert to promote the 1^st^ step or inhibit proper progression to the 2^nd^ step. Since Fyv6, like Slu7 and Prp18, is predicted to interact with Prp8 (**Fig. S1**), we expected that genetic interactions should occur between Fyv6 and Prp8 if the former is also involved in splicing.

**Figure 2.**
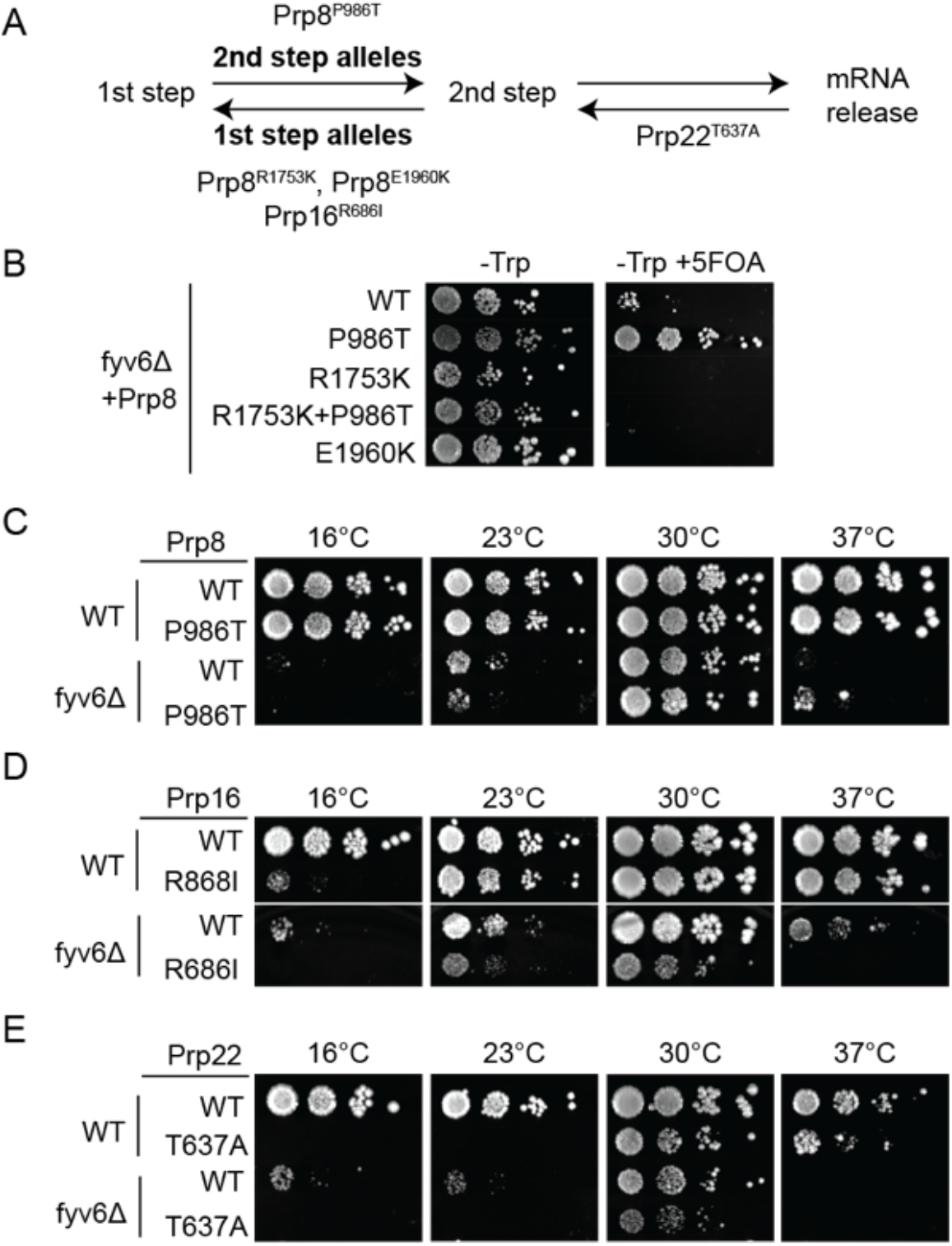
Genetic interactions between Fyv6 and Prp8, Prp16, or Prp22. **A)** Diagram of how Prp8, Prp16, and Prp22 alleles impact the 1^st^ and 2^nd^ steps of splicing. **B)** Alleles of Prp8 were combined with *fyv6Δ* in Prp8 shuffle strains and grown on -Trp or -Trp+5-FOA plates. Yeast growth was imaged after 3 days at 30°C. **C)** Prp8^P986T^/*fyv6Δ* strains were tested for suppression or exacerbation of temperature-dependent growth phenotypes. **D**, **E)** Alleles of Prp16 and Prp22 were combined with *fyv6Δ* and tested for suppression or exacerbation of temperature-dependent growth phenotypes. For panels C-E, yeast were plated on YPD and imaged after 3 (30°C), 4 (23°C and 37°C), or 10 (16°C) days.

Plasmid shuffle of a gene expressing Prp8^E1960K^ into a *fyv6Δ* strain resulted in synthetic lethality even at the normally permissive temperature of 30°C (**Fig. 2B**). Synthetic lethality was also observed for another 1^st^-step allele of Prp8, *prp8-R1753K* (**Fig. 2B**). In contrast, when we shuffled in a 2^nd^-step allele, *prp8-161* (Prp8^P986T^), we observed suppression of the growth defect caused by *fyv6Δ* at both 30 and 37°C (**Figs. 2B**, **C**). When the P986T and R1753K mutations were combined, synthetic lethality was still observed with *fyv6Δ* (**Fig. 2B**). These results are consistent with Fyv6 acting to promote the 2^nd^ step of splicing and its deletion, *fyv6Δ*, promoting the 1^st^ step. Combining *fyv6Δ* with a 1^st^-step Prp8 allele can be synthetically lethal due to failure to properly transition to the 2^nd^ step, while combining *fyv6Δ* with a 2^nd^-step allele may improve yeast growth by facilitating proper 1^st^-step/2^nd^-step equilibrium.

### Genetic Interactions between Fyv6 and Prp16 or Prp22

Prp8 1^st^ and 2^nd^-step alleles, as well as 1^st^ and 2^nd^-step alleles in U6 snRNA and Cef1, can also genetically interact with mutants of the Prp16 or Prp22 ATPases the promote the 1^st^ to 2^nd^-step transition or exit out of the 2^nd^-step by product release, respectively (**Fig. 2A**) (Query and Konarska, 2006, 2012; Eysmont et al., 2019). Based on these observations, we next tested genetic interactions between *fyv6Δ* and Prp16 and Prp22 mutants that presumably slow these conformational changes. Prp16^R686I^ likely slows the 1^st^-to-2^nd^-step transition, leads to a *cs* phenotype at 16°C (Hotz and Schwer, 1998), and is synthetically lethal with 1^st^-step Prp8 alleles (Query and Konarska, 2006). A yeast strain with *fyv6Δ* plus Prp16^R686I^ exacerbates the cold sensitivity, resulting in almost no growth at 16°C and reduced growth at 23°C compared to strains with either allele alone (**Fig. 2D**). The combination of Prp16^R686I^ with *fyv6Δ* also results in reduced growth at 30°C and is synthetic lethal at 37°C. Both *fyv6Δ* and 1^st^-step Prp8 alleles interact negatively with the Prp16^R686I^ ATPase.

The Prp22^T637A^ mutant uncouples ATP hydrolysis from RNA unwinding (Schwer and Meszaros, 2000), likely perturbing the transition out of the 2^nd^-step conformation and product release. Prp22^T637A^ is also a *cs* allele and does not grow at 16 or 23°C (Schwer and Meszaros, 2000 and **Fig. 2E**) and is synthetically lethal with 2^nd^-step alleles of Cef1 (Query and Konarska, 2012). When Prp22^T637A^ and *fyv6Δ* were combined, we did not observe any suppression of the *cs* phenotype of Prp22^T637A^ or the *cs/ts* phenotype of *fyv6Δ* (**Fig. 2E**). Prp22^T637A^/*fyv6Δ* yeast were viable at 30°C but grew more slowly than strains containing only a single allele. Deletion of *FYV6* results in more pronounced genetic interactions with Prp16^R686I^ than with Prp22^T637A^, consistent with the deletion acting as a 1^st^-step allele.

### Impact of fyv6Δ on Yeast Growth using the ACT1-CUP1 Splicing Reporter Assay

If Fyv6 is a component of the splicing machinery as the genetic interactions suggest, we would also predict changes in *in vivo* splicing in the absence of the protein. To test this, we used the ACT1-CUP1 reporter assay in which changes in the splicing of the reporter pre-mRNA (**Fig. 3A**) confer proportional changes in the copper tolerance of a sensitized yeast strain with increased splicing efficiency leading to growth at higher copper concentrations (Lesser and Guthrie 1993). Since yeast lacking Fvy6 grow more slowly than WT even under optimal growth conditions (**Fig. 2**, for example) we scored WT yeast growth on Cu^2+^-containing plates after 48h but *fyv6Δ* yeast were scored after 72h. Consistent with the slow growth and with a function of Fyv6 during splicing, we observed slightly reduced copper tolerance for even the WT ACT1-CUP1 reporter in the *fyv6Δ* strain (**Fig. 3B, C**).

**Figure 3.**
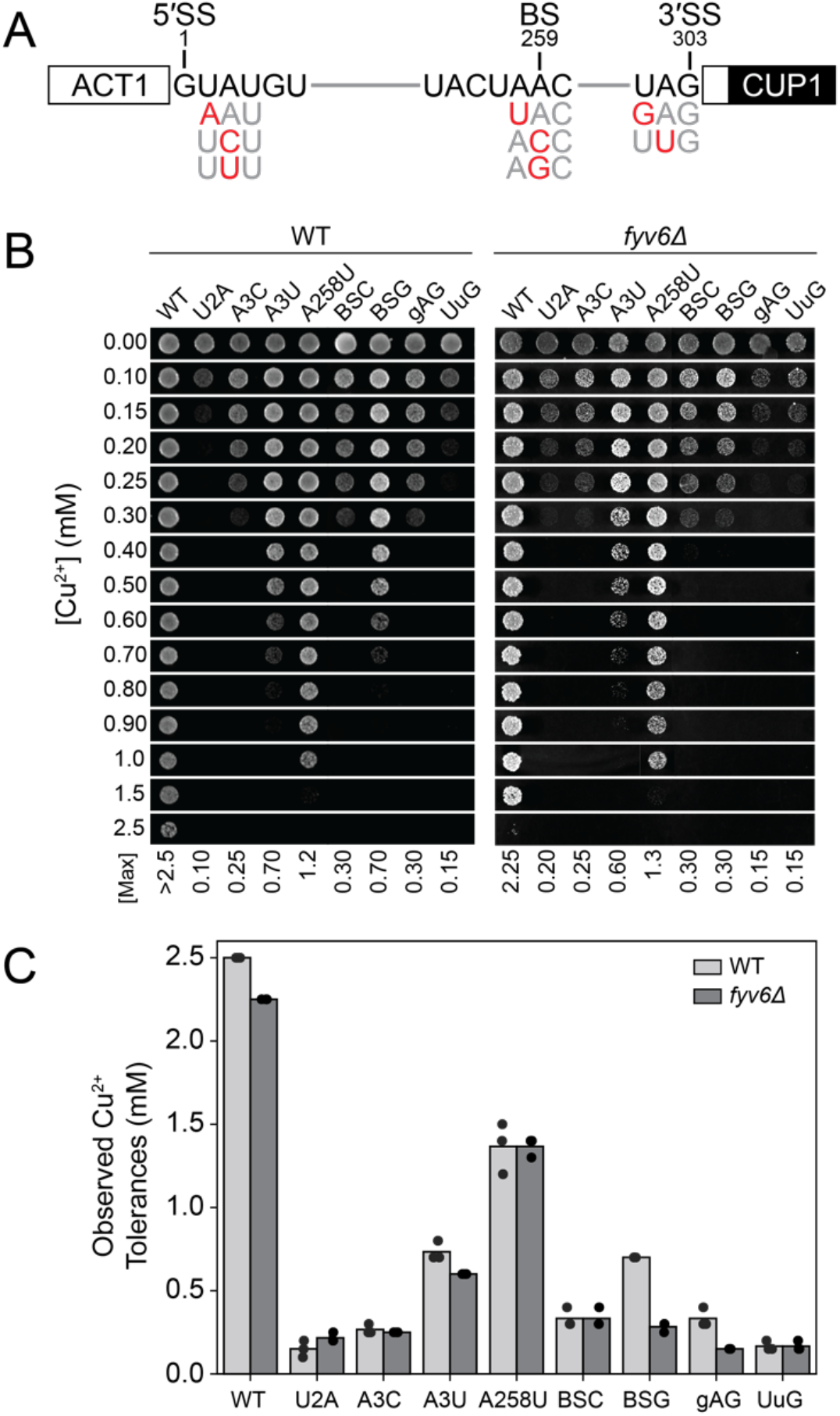
Impact of FYV6 deletion on yeast copper tolerance using the ACT1-CUP1 assay. **A)** Schematic of the WT ACT1-CUP1 reporter along with intronic substitutions. **B)** Images of representative yeast growth on copper containing media shown after 48 (WT) or 72 h (*fyv6Δ*) for strains containing the indicated ACT1-CUP1 reporters. **C)** Maximum copper tolerances observed for each strain for *N=3* replicates (dots). Bars represent the average values.

We also tested several ACT1-CUP1 reporters containing substitutions in the 5’ (U2A, A3C, and A3U) or 3’ SS (UuG and gAG) or the BS (A258U, BSG, and BSC, **Fig 3A**) to determine if loss of Fyv6 is especially detrimental or beneficial to introns with nonconsensus sequences. Copper tolerance was similar between WT and *fyv6Δ* yeast containing reporters with the U2A, A3C, UuG, A258U, and BSC substitutions. However, *fyv6Δ* strains containing A3U, BSG, and gAG reporters exhibited less tolerance to copper than WT (**Fig. 3B, C**), suggesting poorer splicing of pre-mRNAs with these substitutions. Previous work has shown that 1^st^-step alleles of Prp8 result in reduced copper tolerance and few ligated mRNA products with the U2A, A3C, BSC, BSG, UuG, and gAG reporters and that 2^nd^ step alleles of Prp8 or Cef1 improve copper tolerance of yeast with the U2A, BSC, BSG, UuG, and gAG reporters (Liu L. et al., 2007; Query and Konarska, 2012). Several of these reporters are also limiting for the exon ligation reaction itself (U2A, A3C, BSG, UuG, and gAG; Liu L. et al., 2007). Loss of Fyv6 results in changes in copper tolerance most similar to 1^st^-step alleles of Prp8 and is consistent with Fyv6 supporting the 2^nd^ step when present.

### FYV6Δ Changes 3’ SS Selection in an ACT1-CUP1 Splicing Reporter

Finally, since both Prp18 and Slu7 can change 3’ SS selection (Kawashima et al., 2009, 2014; Frank and Guthrie, 1992), we tested whether or not loss of Fyv6 can also change splicing outcomes. We utilized an ACT1-CUP1 reporter containing an additional, alternative 3’ SS proximal to the BS which results in a frameshift when used instead of the distal 3’ SS (**Fig. 4A**). Previous studies with this reporter showed that use of the proximal 3’ SS greatly increases in the presence of the *slu7-1* allele with an ~20-fold change in the ratio of mRNAs produced using the proximal vs. distal sites (Frank and Guthrie, 1992). Indeed, when this reporter was used, we observed the largest differences in copper tolerance (**Fig. 4B**). To confirm that this change in survival was due to a change in 3’ SS usage and use of the proximal SS, we isolated RNAs from the yeast strains and quantified SS usage by primer extension. These results showed an increase in use of the proximal SS and an ~5-fold increase in the ratio of mRNAs produced using the proximal vs. distal sites (**Fig. 4C, D**). Like mutations in Slu7 or loss of Prp18, loss of Fyv6 can also change 3’ SS selection and is consistent with function as a 2^nd^-step splicing factor. These data suggest that Fyv6, like Slu7, helps to enforce a preference for BS distal 3’ SS presumably by facilitating docking of the distal SS and/or preventing docking of the proximal site (Fica et al., 2017).

**Figure 4.**
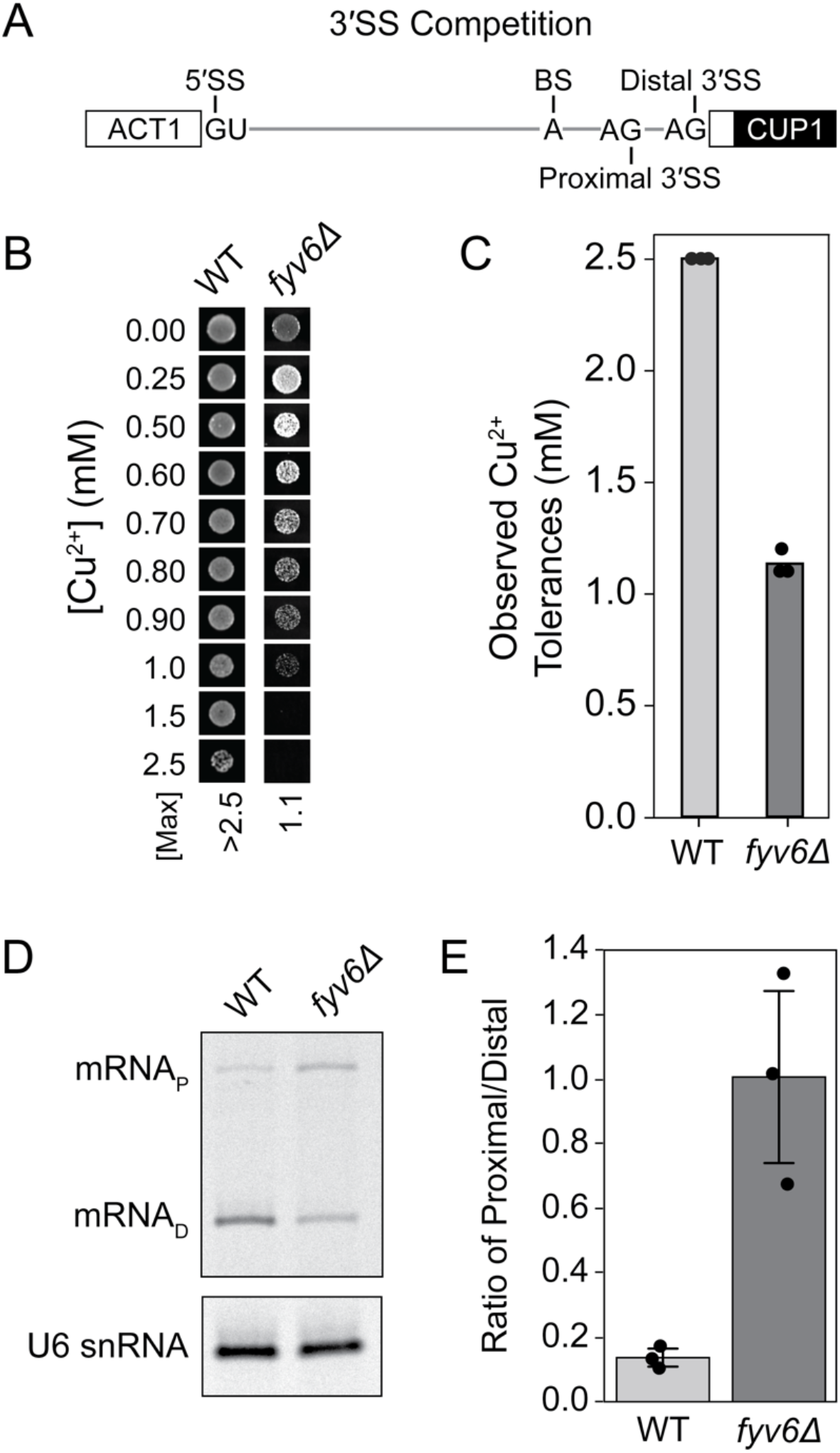
Loss of Fyv6 Changes 3’ SS Selection in Yeast. **A)** Schematic of the 3’ SS competition reporter (3’ SS comp) showing relative locations of the proximal and distal sites. **B)** Images of representative yeast growth on copper containing media shown after 48 (WT) or 72 h (*fyv6Δ*) for strains containing the 3’ SS comp ACT1_CUP1 reporter. **C)** Maximum copper tolerances observed for each strain for *N=3* replicates (dots). Bars represent the average values. **D)** Representative primer extension analysis of mRNAs generated by yeast using the distal (mRNA_D_) or proximal (mRNA_P_) 3’ SS in the presence (WT) or absence of Fyv6 (*fyv6D*). U6 snRNA was analyzed as a loading control. **E)** Quantification of the primer extension results from *N=3* replicates (dots) expressed as a ratio of mRNA_P_/mRNA_D_. Bars represent the average of the replicates ±SD. Means between two experimental groups were compared with an unpaired student’s t-test of equal variance (p = 0.01014).

## Conclusion

Together the genetic and biochemical data presented here as well as the mass spectrometry and cryo-EM work of others indicate that yeast Fyv6 is a 2^nd^-step splicing factor and likely is a component of the spliceosome. Our results do not, however, confirm that the unassigned EM density present in yeast C* and P complex spliceosome cryo-EM maps is due to Fyv6. Further work will be needed to confirm that this is indeed the case either through obtaining higher resolution cryo-EM data or experimental approaches that probe protein-protein interactions to map the Fyv6 binding site.

Many outstanding questions remain about Fyv6 function. We do not know when Fyv6 associates or dissociates from the spliceosome or exactly how it is promoting exon ligation. One possible mechanism could be by direct modulation of Prp22 activity. Several other DExD/H-box ATPases involved in splicing use protein cofactors to stimulate their activity and regulate their association with the spliceosome: Spp2 interacts with the Prp2 ATPase during activation and Ntr1 interacts with the Prp43 ATPase during spliceosome disassembly (Tanaka et al., 2007; Silverman et al., 2004; Warkocki et al., 2015). Fyv6 or human FAM192A could allosterically regulate Prp22 activity by directly interacting with the protein or by helping it be recruited to the correct spliceosomal complex and RNA substrate. Finally, since it seems likely that Fyv6 is the functional yeast homolog of human FAM192A in terms of pre-mRNA splicing, it will be worth investigating if other functions of FAM192A/Fyv6 are conserved in yeast. FAM192A also associates with the 20S proteasome via interaction with PA28γ, a 20S proteasome regulator, for ubiquitin-independent protein degradation within the nucleus (Jonik-Nowak et al., 2018). While yeast lack an apparent homolog for PA28γ (Jonik-Nowak et al., 2018), it may be interesting to determine if Fyv6 is also involved in protein degradation and if there are any Fyv6-dependent links between proteostasis and pre-mRNA splicing.

## SUPPLEMENTAL MATERIAL

Supplemental material is available for this article.

## ACKNOWLEDGEMENTS

We thank Guilaume Chanfreau and his laboratory for helpful discussions and critical review of the manuscript. We thank Peter Ducos and Tim Grant for assistance in making Figure 1.

## METHODS

Yeast strains and plasmids used in this study are described in **Tables S1** and **S2**. Yeast transformation, plasmid shuffling/5-FOA selection, and growth were carried out using standard techniques and media (Treco and Lundblad 1993; Sikorski and Boeke 1991).

### FYV6 deletion

The FYV6 gene was deleted through replacement with a hygromycin resistance cassette by homologous recombination (see **Table S1**; Goldstein and McCusker, 1999). Gene deletion was confirmed by genomic DNA extraction from the strains and PCR amplification of the FYV6 genomic locus using primers Fyv6-check-fwd 5’-TGGATCGAACACAGGACCTC-3’ and Fyv6-check-rev 5’-GTGGAACGAGCAATCAATGTGATC-3’.

### ACT1-CUP1 copper tolerance assays

Yeast strains expressing ACT1-CUP1 reporters were grown to stationary phase in -Leu DO media to maintain selection for plasmids, diluted to OD_600_ = 0.5 in 10% (v/v) glycerol, and spotted onto -Leu DO plates containing 0 to 2.5 mM CuSO_4_ (Lesser and Guthrie 1993; Carrocci et al., 2018). Plates were scored and imaged after 48 h of growth at 30°C for WT strains and after 72 h of growth at 30°C for *fyv6Δ* strains due to differential growth between strains.

### Growth assays

For temperature-dependent growth assays, yeast strains were grown overnight to stationary phase in YPD media, diluted to OD_600_ = 0.5 in 10% (v/v) glycerol, and stamped onto YPD plates. The plates were incubated at 16, 23, 30, or 37°C for the number of days indicated in each figure legend before imaging.

For growth assays in the presence of 5-FOA, yeast strains were grown overnight to stationary phase in -Trp DO media, diluted to OD_600_ = 0.5 in 10% (v/v) glycerol, and stamped onto -Trp and -Trp +5-FOA plates. The plates were incubated at 30°C for 3 days before imaging.

### Primer extension

Cell cultures were inoculated from stationary phase saturated cultures grown overnight in -Leu DO media. The cultures were then grown until OD_600_ = 0.6 −0.8, and 10 OD_600_ units were collected by centrifugation. Total RNA was isolated from yeast and contaminating DNA was depleted using the MasterPure Yeast RNA Purification Kit (Epicentre, Madison, WI) protocol with minor changes as previously described (Carrocci et al., 2017). IR700 dye conjugated probes (Integrated DNA Technologies, Skokie, IL) were used for primer extension of the ACT1-CUP1 reporter (10 pmol yAC6: /5IRD700/GGCACTCATGACCTTC) and U6 snRNA (2 pmol yU6: /5IRD700/GAACTGCTGATCATGTCTG) (Carrocci et al., 2017; van der Feltz et al., 2021). Primer extension products were visualized on a 7% (w/v) denaturing polyacrylamide gel (42 cm x 22 cm x 0.75 mm) run at 35W for 80 min at RT. Gels were imaged with an Amersham Typhoon NIR laser scanner (Cytiva), and band intensities were quantified with Image J (version 1.53v, 2022).

### Network analysis of potential Fyv6 interactions

Protein-protein and protein-RNA interactions found in the atomic model for the P complex spliceosome (6BK8) were identified using LouiseNET and the resulting nodes and edges were plotted as an undirected network model using GEPHI as previously described (Bastian et al., 2009; Kaur et al., 2022).

